# BDNF Val66Met polymorphism as putative genetic substrate of music-induced plasticity in auditory prediction

**DOI:** 10.1101/2021.04.07.438769

**Authors:** S.E.P. Bruzzone, L. Bonetti, T. Paunio, K. Kantojärvi, M. Kliuchko, P. Vuust, E. Brattico

## Abstract

Predictive processing of sounds depends on the constant updating of priors based on exposure to posteriors, which through repeated exposure mediates learning. The result of such corrections to the model is seen in musicians, whose lifelong training results in measurable plasticity of audio-motor brain anatomy and functionality. It has been suggested that the plasticity of auditory predictive processes depends on the interaction between the environment and the individual’s genetic substrate. However, empirical evidence to this is still missing. BDNF is a critical genetic factor affecting learning and plasticity, and its widely studied functional variant Val66Met single-nucleotide polymorphism offers a unique opportunity to investigate neuroplastic functional changes occurring upon a years-long training. We hypothesised that BDNF gene variations would be driving neuroplasticity of the auditory cortex in musically trained human participants. To this goal, musicians and non-musicians were recruited and divided in Val/Val and Met carriers and their brain activity measured with magnetoencephalography (MEG) while they listened to a regular auditory sequence containing different types of prediction errors. The auditory cortex responses to prediction errors was enhanced in Val/Val carriers who underwent intensive musical training, compared to Met and non-musicians. Our results point at a role of gene-regulated neurotrophic factors in the neural adaptations of auditory processing after long-term training.

## Introduction

According to the predictive coding theory (PCT), we constantly anticipate the statistical regularities of the environmental stimuli based on internal models or priors, that are automatically updated according to the domain-specific experience (Friston, 2012; Koelsch et al., 2019; Vuust et al., 2009). This predictive process occurs within the sensory cortex, which, after a repeated presentation of stimuli having specific features, rewires itself and responds more vigorously to errors of predictions. The ability to predict the outcomes based on previously learned information is fundamental to adapt to the continuously evolving environmental demands and is the result of a complex interplay between genetic factors, physiological and cognitive processes (Bramham and Messaoudi, 2005; Z. J. Huang et al., 1999; Irvine, 2018; Keuroghlian and Knudsen, 2007).

In the auditory cortex, updates occurring upon violations to the auditory models are reflected by increased neuronal activity in supratemporal and fronto-parietal areas, as indexed in two subsequent event related potentials/fields (ERP/ERF), namely the mismatch negativity (MMN) and the P3a. Occurring within 100-200ms from the deviant stimulus (Näätänen et al., 1978, 2007), the MMN has been linked to the bottom-up process of automatic error detection (Harms et al., 2020; Vuust et al., 2009) while the later P3a, occurring around 250-300ms (Escera and Corral, 2007; Ruusuvirta et al., 1995; Sams et al., 1983), is thought to mirror the top-down process of evaluation of the error and updating of the model and is also a highly hereditable tract of brain cognitive processes (O’Connor et al., 1994).

The continuous learning process updating the accuracy of auditory predictions is well exemplified by professional musicians, whose auditory predictive abilities are enhanced as a result of their years-long music training (Brattico et al., 2001, 2009; Herdener et al., 2010; Jäncke, 2009; Kliuchko et al., 2019a; Kraus & Chandrasekaran, 2010; Vuust et al., 2012b). Musicians not only display changes in motor functions related to their playing practice (Kraus and Chandrasekaran, 2010; Matrone and Brattico, 2021; Reybrouck and Brattico, 2015; Reybrouck et al., 2018; Schlaug, 2015), but also their auditory perception is more finely tuned to differences in sound features than those of non-musicians, as indexed by their stronger MMN in response to metric and melodic violations (Vuust et al., 2005, 2009).

Training-dependent neurophysiological changes have been proposed to depend on a combination of environmental and genetic factors (Bengtsson et al., 2005; Imfeld et al., 2009; Steele et al., 2013; Steele & Zatorre, 2018). Indeed, genetic basis of musical competence is under exploration with promising initial findings (Mosing et al., 2014,2016; Pulli et al., 2008; Ukkola et al., 2009). Based on evidence of association between music-related behavior and familial factors, some scholars have proposed that music-induced neuroplasticity might result from the individual genetic repertoire (Wesseldijk et al., 2019,2020). However, this relationship has yet to be confirmed empirically (Merrett et al., 2013; Steele and Zatorre, 2018).

In this framework, the Val66Met, a common single-nucleotide polymorphism (SNP) occurring in the gene of the brain-derived neurotrophic factor (BDNF), offers a unique opportunity to clarify the genetic implications of training-dependent plasticity. BDNF is a widespread neurotrophin that plays a key role in neural development and brain plasticity, mediating neuronal differentiation, synaptogenesis, long-term potentiation (LTP) and depression (LTD) (Binder and Scharfman, 2004; Huang and Reichardt, 2001; Poo, 2001; Tyler and Pozzo-Miller, 2001), with a significant impact on cognitive processes underlying learning and memory. Notably, BDNF also promotes the development and the later adaptations of the auditory cortex, where it mediates the refining of the auditory cortical receptive fields based on environmental demands (Chang et al., 2005; Chaudhury et al., 2013; Guo et al., 2013; Kaur et al., 2004; Singer et al., 2014).

Research on animal models reports an association with low levels of BDNF and morphological abnormalities in the hippocampal neurons (Messaoudi et al., 2002) as well as to impairments in NMDA-mediated transmission and LTP processes (Caldeira et al., 2007; Ninan et al., 2010; Pattwell et al., 2012). Conversely, increased BDNF levels in hippocampus are accompanied by enhanced neurogenesis and plasticity upon the exposure to auditory enriched stimulation in the form of music (Angelucci, Fiore, et al., 2007; Chikahisa et al., 2006; Xing et al., 2016; for a review, see Brattico et al., 2021).

Similarly, in humans, greater BDNF plasma levels have been found after listening to music (Nair et al., 2020) and in musicians compared to non-musicians (Minutillo et al., 2020; Oikkonen et al., 2016), reflecting environmental-dependent physiological changes. Moreover, genetics-dependent high baseline levels of BDNF have been associated to increased plasticity and better cognitive performances (Egan et al., 2003; Gratacòs et al., 2007; Hariri et al., 2003; Ho et al., 2006; Lu et al., 2015; Pezawas et al., 2004). Specifically, the availability of circulating BDNF can be affected by the Val66Met (or rs6265) SNP, a common missense mutation that involves the substitution of a valine (Val) with a methionine (Met) in position 66 of the pro-domain region of the BDNF precursor, which is indispensable for the trafficking and secretion of the mature peptide. When mutated, the secretion of BDNF is reduced, causing individuals carrying the Met allele having lower baseline levels compared to Val carriers (Egan et al., 2003; Hariri et al., 2003; Lang et al., 2009; Seidaha et al., 1996).

In support of this, the lower BDNF levels found in Met carriers have been linked to altered episodic memory, brain structure and function in both healthy (Egan et al., 2003; Lu et al., 2015) and clinical populations (Gratacòs et al., 2007; Hariri et al., 2003; Ho et al., 2006; Pezawas et al., 2004; Spalletta et al., 2010; Szeszko et al., 2005; Wang et al., 2014). Evidence from magnetic resonance imaging (MRI) studies reports reduced volume in hippocampus (Bueller et al., 2006; Pezawas et al., 2004; Schofield et al., 2010; Wang et al., 2014), prefrontal cortex (Pezawas et al., 2004), and temporal and occipital lobes (Ho et al., 2006) for Met carriers (Val/Met and Met/Met carriers) compared to Val/Val carriers. Further, several studies described anomalous hippocampal activity during memory tasks associated to the Met allele (Egan et al., 2003; Hariri et al., 2003; Schofield et al., 2010). Consistent with this, a study by Schofield et al. (2009) reported reduced P3b responses to an oddball paradigm in Met carriers, as a reflection of hippocampal abnormal activity in Met carriers.

Altogether, these findings support the influence of BDNF Val66Met on hippocampus-dependent memory and cognitive processes, in line with the key role exerted by BDNF at the molecular level. Moreover, the influence of BDNF on glutamatergic transmission, which is crucial for the generation of MMN (Harms et al., 2020; Javitt et al., 1996; Parras et al., 2020; Tikhonravov et al., 2008), and the involvement of the hippocampus in novelty processing during oddball tasks (Crottaz-Herbette et al., 2005; Halgren et al., 1995a; Halgren et al., 1995b), suggest that differential levels of the neurotrophin could affect the MMN generation. This evidence, together with the implication of BDNF in the development and retuning of the auditory cortex, point to a possible influence of Val66Met in learning processes mediated by auditory predictions and, specifically, that such processes could be particularly enhanced upon long-term musical training.

Nonetheless, although the role of Val66Met SNP in cognition has captured the attention of a discrete amount of studies, evidence on the contribution of the polymorphism to the neurophysiology of auditory predictive processes is still largely missing. However, in a recent work by Bonetti et al., (2021), another polymorphism, the Val158Met SNP in the catechol-O-methyl-transferase (COMT) gene, has been shown to affect auditory predictive processes, suggesting a genetic influence on the generation of MMN. Hence, in the present work we aim to shed light on the genetic contribution to experience-dependent, particularly music-induced plasticity as indexed by the magnetic counterparts of MMN and P3a, namely MMNm and P3am. Given the pivotal role played by BDNF in neural development and plasticity, we hypothesize that the neuroplastic changes induced by intensive musical training would be modulated by different variants of the BDNF Val66Met polymorphism. Specifically, we expect the brain responses to environmental irregularities to be enhanced in musicians carrying the Val/Val allele compared to musicians carrying the Met allele. For the first time to our knowledge, we investigate the modulatory effects of BDNF on cortical plasticity in auditory predictive processes resulting from years-long musical training. For this purpose, using magnetoencephalography (MEG), we measured the neurophysiological activity of musicians and non-musicians while they were passively presented with repetitive and error sounds. Next, we used a combination of magnetoencephalography (MEG) and MRI to detect and localize the sources of neurophysiological activity focusing on the prediction error signals originating from the auditory cortex.

## Results

### Genetic and demographic data

The distribution of the BDNF genetic variant in our sample was: Val/Val = 57 (77.03%); Val/Met = 16 (21.62%) and Met/Met = 1 (1.35%), coherent with the Hardy–Weinberg equilibrium (p-value < 1×10-6).

Detailed demographic information is reported in **Table 1**.

**Table 1.** Demographic data of the participants described according to their BDNF genetic variation (M1 = musicians Val/Val; NM1 = non-musicians Val/Val; M0 = musicians Val/Met and Met/Met; NM0 = non-musicians Val/Met and Met/Met). The table shows means for age (± indicates standard deviation) and frequencies for number of participants, sex (M = males; F = females) and handedness (L = left-handed; R = right-handed; A = ambidextrous).

| Information | Whole sample | M1 | M0 | NM1 | NM0 |
| --- | --- | --- | --- | --- | --- |
| *Number of participants* | 74 | 19 | 8 | 38 | 9 |
| *Age* | 28.70 ± 8.34 | 29.58 ± 7.21 | 28.38 ± 9.41 | 28.16 ± 9.11 | 29.44 ± 7.21 |
| *Sex* | M = 32; F = 42 | M = 8; F = 11 | M = 5; F = 3 | M = 17; F = 21 | M = 2; F = 7 |
| *Handedness* | L = 5; R = 67; A = 1 (*1 missing*) | L = 2; R = 16 (*1 missing*) | L = 0; R = 7; A = 1 | L = 2; R = 36 | L = 1; R = 8 |
| *Years of formal music training* | 6.56 ± 9.03 | 16.00 ± 4.74 | 19.62 ± 10.24 | 0.59 ± 1.05 | 0.22 ± 0.66 |

## BDNF and neural responses to deviants

### MEG sensors

First, we computed the neural responses to deviants to visually inspect the quality of the data and to control the clarity of the isolated MMN and P3a peaks. Coherently with the previous literature that used the same paradigm as the one used in this study (Bonetti et al., 2017, 2018, 2021; Kliuchko et al. 2016, 2019a), we assessed that such peaks were stronger for musicians compared to non-musicians. In agreement with what reported in those studies, we observed that the clearest MMN and P3a peaks occurred for localization, slide and timbre deviants, while rhythm, intensity and pitch showed a reduced response. Waveforms, topoplots and neural sources are depicted in **Figure 1** and reported in details in **Table ST1**. Second, we used ANOVAs to compare the neural responses to deviants of our four experimental groups (M1, M0, NM1, NM0), focusing especially on the post-hoc analysis between M1 and M0. This analysis was computed for each combined planar gradiometer and each time-point within the time-range 0 – 400ms. Then, they were corrected for multiple comparisons using the Tukey-Kramer post-hoc and a cluster-based Monte-Carlo simulation (see Methods for details and **Figure 2** for a graphical illustration). In particular, **Table 2** and **Figure 3A** show the significant clusters where the ANOVAs were significant and M1 was stronger than M0 (and vice versa). Our results indicated that musicians had overall stronger neural responses to deviants when carrying the BDNF allele Val/Val than carrying the Val/Met and Met/Met alleles. In the latter case, the neural responses to deviants appeared very similar to the non-musicians ones. Notably, the largest clusters were observed for localization (k = 79; p < .001), slide (k = 53; p <.001) and timbre (k = 36; p < .001), which correspond to the deviants that enhanced the clearest MMN peaks (**Figure 3A** and **3B**). Conversely, the deviants that enhanced a less clear MMN (rhythm, pitch and intensity) returned smaller and less defined clusters.

**Figure 1.**
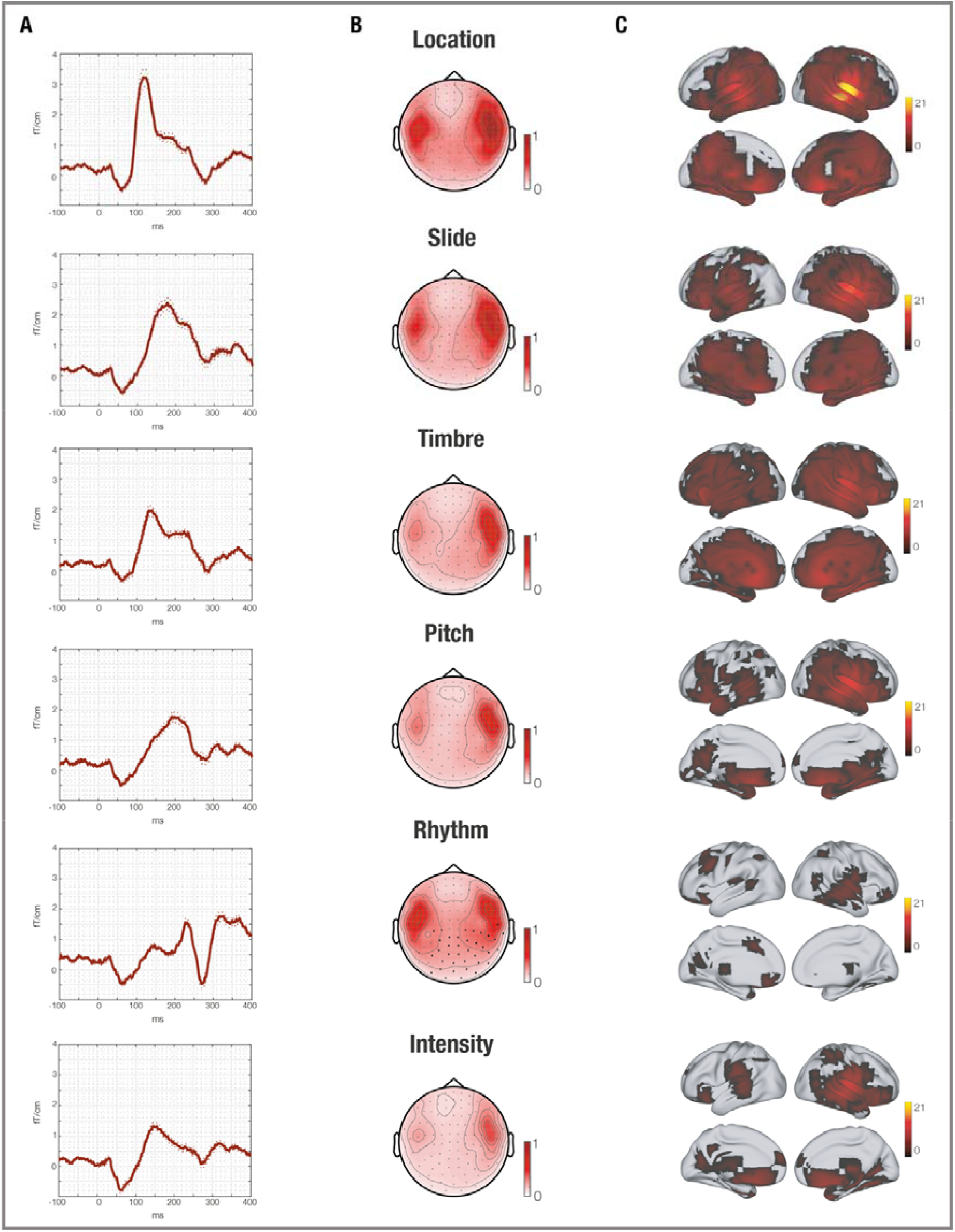
Neural responses to deviants. Waveforms (A), topoplots (B) and brain sources (C) of the neural signal responses to deviants, shown for all participants and separately for musicians and non-musicians. Specifically, waveform images refer to the channel where we recorded the peak amplitude in response to deviants. The topoplots (fT/cm) and neural sources indicate the location of the brain activity at the peak amplitude depicted in the waveforms. In the brain templates, we depicted the t-values emerging from the contrast between neural responses to deviants and baseline.

**Figure 2.**
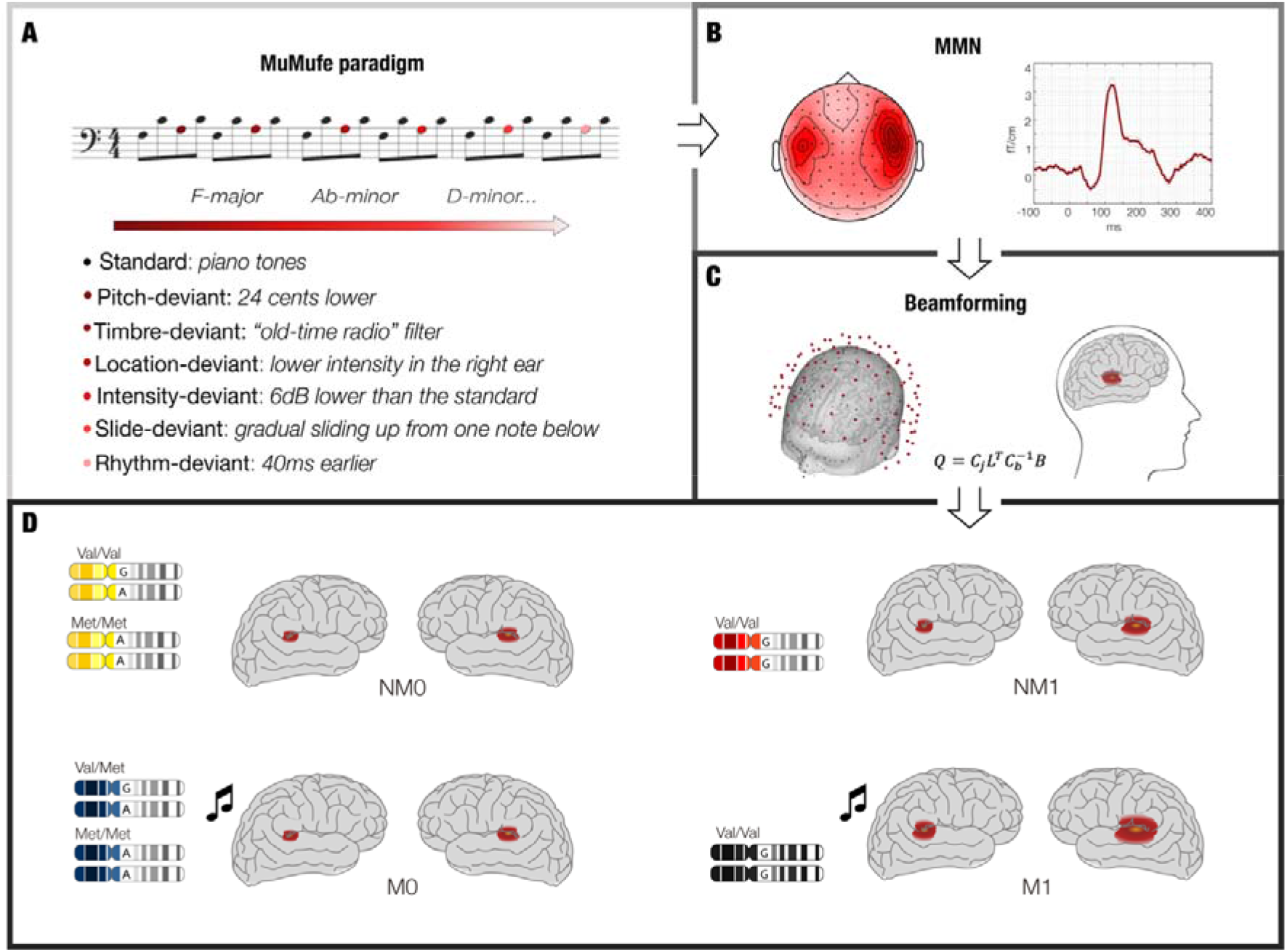
Overview of the analysis pipeline. **A** – Participants were presented with the musical multifeatures paradigm (MuMufe) during MEG recording. With this paradigm, we obtained neural responses to deviant acoustic stimulations. **B** - MEG data has been collected and pre-processed and typical event-related components elicited by deviant stimulation such as mismatch negativity (MMN) have been identified. **C** – Source reconstruction has been estimated by using an MEG overlapping sphere forward model and a beamforming approach as inverse model. **D** – Participants have been divided into different groups according to their BDNF genetic variation and their musical expertise and their different brain response to deviants tested through analysis of variance (ANOVA). Specifically, we split the participants into four groups: musicians Val/Val (i – M1); musicians Val/Met and Met/Met (ii – M0); non-musicians Val/Val (iii – NM1); non-musicians Val/Met and Met/Met (iv – NM0).

**Table 2.** Significant clusters outputted by MCS (p < .001) on the significant results provided by the ANOVAs contrasting the four experimental groups (M1, M0, NM1, NM0) for each time-point. Specifically, the table depicts the contrasts between the neural responses to deviants of M1 vs M0, highlighting the higher neural responses recorded for musicians BDNF Val/Val vs musicians BDNF Val/Met and Met/Met. The analysis has been conducted independently for each of the six deviants (see Methods for details). k refers to the spatio-temporal extent of the cluster (e.g., the overall number of channels and time-points forming the cluster).

| Deviant | k M1 > M0 | k M0 > M1 |
| --- | --- | --- |
| *Localization* | 79 | 43 |
|  | 8 | 10 |
|  | 8 | - |
|  | 8 | - |
|  | 8 | - |
| *Slide* | 53 | - |
|  | 9 | - |
| *Timbre* | 36 | - |
|  | 16 | - |
|  | 8 | - |
| *Pitch* | 14 | 12 |
|  | 6 | 8 |
| *Rhythm* | 24 | 16 |
|  | 8 | - |
| *Intensity* | - | 4 |

**Figure 3.**
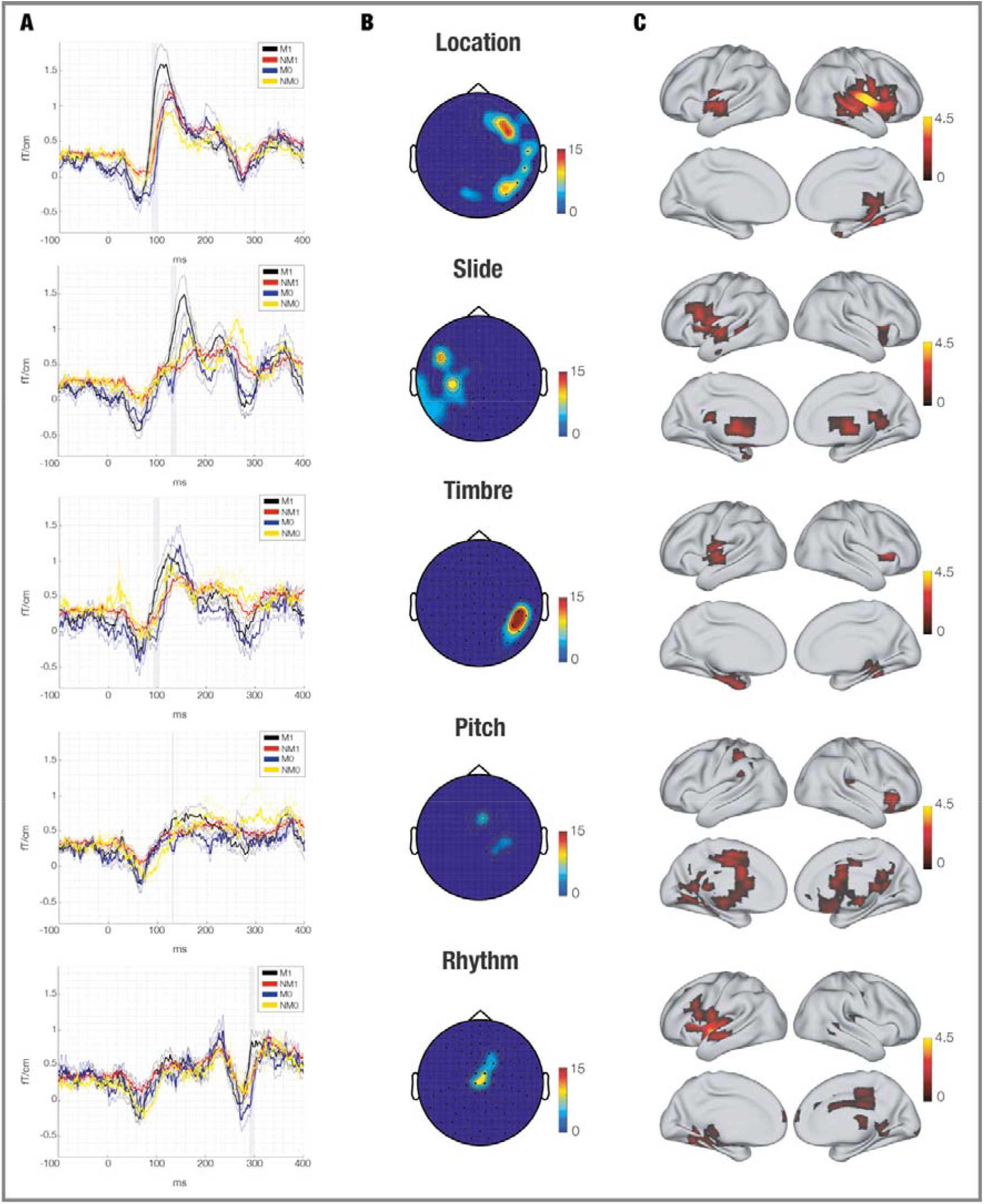
Neural responses to deviants illustrated according to the BDNF polymorphism groups. Waveforms (A), topoplots (B) and brain sources (C) of the neural signal recorded in response to deviant stimulations. Grey areas in the waveforms show the significant time-points emerged by the MCS performed on the results of the ANOVAs contrasting the four experimental groups (M1, M0, NM1, NM0). The topoplots illustrates the temporal extent (in ms) and the localization of the MEG channels that were significantly different after computing MCS on the ANOVAs results. Brain sources depict the t-values of the contrast between M1 and M0 computed in the significant time-points emerged by the ANOVAs on MEG sensors. This analysis allowed us to identify the specific brain regions where BDNF Val/Val musicians presented stronger responses to deviants than BDNF Val/Met and Met/Met musicians.

### Source localized activity

The brain sources of the MEG signal were estimated in the significant time-windows that emerged from the MEG sensor analysis. For each source-reconstructed brain voxel and time-point, we used a beamforming approach in combination with general linear models (GLM). This analysis was computed independently for each participant. Next, we computed two group-level analyses. In the first case, we did not contrast any different group of participants. Instead, we merely assessed the quality of the source reconstruction, to make sure that the MMN and P3a were properly localized in source space (**Figure 1C**). After this, we computed a second group-level analysis contrasting M1 vs M0, aiming to isolate the brain areas where M1 reported the strongest brain activity. This analysis was conducted based on the significant time-points emerged from the analysis on MEG sensors. Multiple comparisons were corrected by employing a cluster-based permutation test (Hunt et al., 2012; see Methods for details).

As described in **Table ST2**, this analysis revealed stronger brain activity for M1 than M0, mainly localized in the temporal brain areas associated to the MMN generation, such as insula and superior temporal gyrus. As illustrated in **Figure 3C**, the analysis returned the significant clusters for the following deviants: (in this case k refers to the number of voxels forming the significant clusters): localization (k = 115; p < .05), slide (k = 63; p < .05), timbre (k = 36; p < .05), pitch (k = 51; p < .05) and rhythm (k = 80; p < .05).

## Discussion

In the present study we investigated the genetic contribution of BDNF Val66Met SNP to training-dependent plasticity in auditory predictive processes, as indexed by MMNm. Specifically, we found that Val/Val carriers who underwent years-long musical training had overall enhanced neurophysiological prediction error signals compared to both their musician and genetic counterparts, as a result of neuroplastic changes in their neural predictions. By means of a relatively complex auditory stimulation, MMN were elicited for six different types of deviants, with the strongest MMN error signal in response to changes in localization, followed by those in slide and timbre. The weakest MMN signal was found for intensity, consistent with previous studies using the same auditory paradigm and a subset of these participants (Bonetti et al., 2017, Bonetti et al., 2018; Bonetti et al., 2021; Kliuchko et al., 2016; Vuust et al., 2011; Vuust et al., 2012b). As expected, source reconstruction analysis of the MEG sensors showed stronger neural activation within the temporal areas associated with MMN generation, proportionally to the intensity of the signal generated by the single deviants.

Consistent with former findings about experience-dependent plasticity in musicians, we found enhanced auditory ERPs in musicians compared to non-musicians, with greater magnitude of both MMN components (Herholz and Zatorre, 2012; Kraus and Chandrasekaran, 2010; Lappe et al., 2008; Shahin et al., 2003; Vuust et al., 2011, 2012b, 2012a). Nonetheless, since the main focus of the study was on the role played by BDNF in neuroplastic changes induced by training, and since the differences between musicians and non-musicians have been widely described by former studies (Brattico et al., 2001; Kliuchko et al., 2019b; Schlaug, 2015; Steele et al., 2013; Vuust et al., 2005; Vuust et al., 2012b) we did not focus on this detail. Instead, we mainly focused on the differences between musicians in relation to their genotype.

In accordance with our hypothesis, we observed an effect of the Val66Met polymorphism on the MMN of Val/Val carriers who received musical training. Specifically, we found main differences for consecutive timepoints occurring within the window of MMN generation. Conformingly to the differential strengths of response across the deviants, the M1 vs M0 contrast revealed the largest significant clusters for location. Although the deviant clusters were generally modest in size, between-group differences were consistently found within 100-200ms from the onset of the deviant stimulus, with the sole exception of rhythm, for which the difference was greatest around 300ms. Nonetheless, previous studies using the same paradigm as the one used here (Bonetti et al., 2017; Kliuchko et al., 2019b) reported a later MMN component for the rhythm deviant compared to the other musical features. Thus, it is plausible to assume that the observed modulatory effect of the polymorphism affected a subset of the physiological processes involved in MMN generation. Moreover, the relatively stringent statistical analyses used to identify the relevant time points support the reliability of such significance. Conversely, no significant differences were found in the characteristic time window for P3a.

Contrasts of the reconstructed sources revealed stronger brain activity for the M1 group vs M0 in insula, Heschl’s gyrus, frontal (or Rolandic) operculum and superior frontal gyrus occurring within the time window of the MMN for all deviants. The difference between the two groups was greater for localization, slide, timbre, and rhythm, with the strongest effects for localization. Again, the degree of brain activation was correlated to the Val homozygous group and was stronger for MMNs elicited by localization, slide, and timbre deviants than those elicited by pitch and intensity. Notably, the contrast revealed a significant higher activation of the auditory areas for the M1 group also in response to rhythm. In addition, minor sources were reconstructed also in the hippocampus and in the parahippocampal area.

Coherently with our results, previous literature showed differential morphology of temporal and insular areas that have been previously associated to the Val66Met SNP. In an MRI study conducted on a large cohort of healthy volunteers, Wang et al. (2004) reported increased surface expansion of the anterior insular cortex, accompanied by a greater connectivity with the dorsolateral prefrontal cortex (DLPFC) for Val/Val carriers as compared to Met carriers. Similarly, Ho et al. (2006) reported reduced grey matter in the left temporal and superior frontal gyrus of healthy Met carriers. Finally, multiple studies reported associations between the Met allele and reduced hippocampal volume accompanied by abnormal activity (Bueller et al., 2006; Egan et al., 2003; Hariri et al., 2003; Pezawas et al., 2004; Schofield et al., 2010; Wang et al., 2014). Although we focused on functional activity and not on volumetric differences, the significant brain areas reported in our study largely overlap with the areas previously described as Val66Met endophenotypes. In this framework, functional changes might reflect different structural features that depend on genotype.

The results of our study, together with the previous findings on BDNF Val66Met, indicate that this SNP orchestrates a subtle yet consistent modulation in the structure and function of temporal areas. As BDNF plays a critical role in cortical development, it is not unlikely that, being exposed to low BDNF baseline levels since birth, Met carriers present such anatomical differences. Nonetheless, MMN responses of Val/Val non-musicians were not significantly enhanced compared to those of Met/Met and Val/Met non-musicians. Instead, the difference emerges clearly when the two groups of musicians are compared. Thus, because BDNF is a mediator of synaptic plasticity throughout life, the effects of the Val66Met polymorphism might appear only upon year-long activities that directly involve BDNF signalling, such as extensive practice of a musical instrument. Moreover, the stronger MMN amplitude in M1 group suggests that the higher BDNF concentrations of the Val/Val genotype may favour the development of a better functional optimization of the auditory cortex after a long-term training. Indeed, expressed and released by cortical pyramidal neurons (Conner et al., 1997), BDNF promotes the functional organization of the auditory cortex by regulating the development of parvalbumin inhibitory interneurons. In the auditory cortex, inhibitory neurons regulate experience-dependent adaptations, by sharpening the receptive fields of the auditory neurons and promoting their maturation (Chang et al., 2005; Froemke et al., 2007; Guo et al., 2013; Kaur et al., 2004). Thus, BDNF promotes functional and perceptual long-lasting changes to auditory stimuli (Singer et al., 2014), being one of the key factors underlying the plastic changes favouring the augmented ability to discriminate sound features which is proper of musicians.

This claim is also supported by the higher BDNF plasma levels found in musicians compared to non-musicians (Minutillo et al., 2020), which might reflect the enhanced plastic properties reported by previous neuroimaging and neurophysiology studies (Bengtsson et al., 2005; Imfeld et al., 2009; Kraus and Chandrasekaran, 2010; Schlaug, 2015; Steele et al., 2013; Vuust et al., 2012b, 2012a). Notably, in this study we demonstrated that such plastic properties are influenced by the Val66Met polymorphism.

Furthermore, the enhanced MMN responses found in the M1 group might be explained in consideration to the role of BDNF in regulating glutamatergic transmission and NMDA-dependent LTP (Bramham and Messaoudi, 2005; Messaoudi et al., 2002; Ninan et al., 2010; Pattwell et al., 2012). Thus, because the generation of MMN requires NMDA transmission, higher BDNF levels might indeed facilitate the communication and plasticity of cortical pyramidal neurons, resulting in greater amplitude of the signal recorded at the scalp for M1 vs M0.

Altogether, our findings revealed a significant relationship between Val66Met and experience-dependent plasticity, showing and enhanced neuroplasticity of Val/Val carriers upon long-lasting musical training, as indexed by the stronger MMN elicited in Val/Val musicians compared to Met carrier musicians.

### Limitations and future perspectives

The present study aimed to investigate the role of BDNF Val66Met polymorphism on experience-dependent plasticity in musicians. A limitation of this study is represented by the low statistical power resulting from the small experimental populations examined. In particular, the Met carriers (N=17) were only seventeen whereas the Val/Val carriers were 57, with a consistent difference between the two non-musician groups (M1: N=19, M0: N=38). In turn, the musician groups were of comparable size (M1: N=9, M0: N=8), but their absolute numerosity was scarce. Moreover, only one subject was homozygous for the Met allele, which did not allow us to investigate the differences according to the three possible BDNF variants independently. Nonetheless, this is in line with the distribution of the Met/Met allele across the population as well as with the previous studies about Val66Met (Gratacòs et al., 2007; Hariri et al., 2003; Ho et al., 2006; Kleim et al., 2006; Pezawas et al., 2004; Wang et al., 2014).

Furthermore, the relatively small number of participants in M0 and M1 might limit the reliability of the reconstruction of the MEG sources. While the insular cortex is superficial enough to have its activity reliably detected by MEG, the estimate of activity from subcortical regions is generally less precise. Nonetheless, given the involvement of the hippocampus in oddball paradigms (Sonia Crottaz-Herbette et al., 2005; Halgren et al., 1995a; Halgren et al., 1995b) and the previous evidence regarding hippocampal impairments in relation to the Met allele (Egan et al., 2003; Hariri et al., 2003; Schofield et al., 2010), it cannot be excluded that the reduced MMN responses in Met carriers are also linked to this area.

Finally, it is important to underline the correlational nature of the results presented in this study. Given the complexity of the cognitive processes involved in prediction error and learning, it is plausible to assume the genetic contribution to these functions depends not only on a single SNP but on the combination of a larger pool of genes, together with their diverse interaction with the environment. Although our study provided a first picture of the relationship linking genetics and experience-dependent neurophysiological changes in humans, further research is called to shed light on the influence exerted by such genetic and environmental factors on neural plasticity. On the genetic level, dissociation studies considering candidate genes that are likely not to be implicated in these processes might help clarify the centrality of BDNF polymorphisms, while on the environmental level a better control on the total hours of practice and the age of its beginning would provide greater insights on the role of BDNF in learning.

## Materials and methods

### Participants

The participants were part of the broader project “Tunteet”, a research protocol involving the collection of neurophysiological, genetic, behavioural and psychological measures regarding cognitive and musical skills, audition and affective behaviour. The complete dataset consisted of 140 volunteers, recruited in the area of Helsinki, Finland. The protocol was approved by the Coordinating Ethics Committee of the Hospital District of Helsinki and Uusimaa (approval number: 315/13/03/00/11, obtained on March the 11th, 2012). The procedures were conducted complying with the ethical principles stated by the Declaration of Helsinki. Further information about this research protocol and related findings are reported in separate studies (Alluri et al., 2017; Bonetti et al., 2017, Bonetti et al., 2018, Bonetti et al., 2021; Burunat et al., 2015; Burunat et al. 2016; Criscuolo et al., 2019; Haumann et al., 2021; Kliuchko et al. 2015, 2016, 2018, 2019; Saari et al., 2018).

All participants were in good health, had normal hearing and were not under any medication. Further, no history of neurological nor psychiatric disease was reported. An informed consent was signed before the beginning of the experiment and a compensation for the participation in the study was given through vouchers that they could use for culture and sports (e.g., concerts, museums or swimming pools.

Participants were divided into musicians (M) and non-musicians (NM) according to the criteria previously used by Criscuolo et al. (2019). The information on their musical expertise was obtained by combining data from a paper and pencil questionnaire (as successfully used in previous works: e.g., Brattico et al., 2009, Criscuolo et al. (2019)) as well as an online survey named Helsinki Inventory for Music and Affect Behavior (HIMAB) (Gold et al., 2013). On the basis of such information, participants were classified as ‘musicians’ when they met the following criteria: (i) having more than 5 years of music practice; (ii) considering themselves as musicians; (iii) having obtained a final degree at a music academy, being involved in musical education or monetarily compensation for their music performance or teaching activities. Conversely, participants with less than three years of musical training were considered as ‘non-musicians’.

Then, the two groups were furtherly classified according to their genotype: group 1 included Val/Val carriers, whereas group 0 Met/Met and Met/Val (0) carriers. Due to the low frequency of Met/Met homozygotes, that would lead to a poor statistical power, Val/Met and Met/Met genotypes were merged into one single group for all analyses. Sixty-six participants were excluded for missing genetic information or because they did not encounter the criteria needed for the subdivision between musicians and non-musicians. Detailed information about demographic data and musical expertise for the final pool of participants (n = 74) is reported in **Table 1**.

### Experimental design and stimuli

Mismatch negativity (MMN) responses were evoked with a fast musical-multifeature MMN paradigm (MuMuFe), involving four tones patterns arranged in an “Alberti bass” configuration (Vuust et al., 2012a; Bonetti et al., 2017; Bonetti et al., 2021; Kliuchko et al., 2016; Vuust et al., 2011). Sound stimuli were synthesized with the software sampler “Halion” in Cubase (Steinberg Media Technologies GmbH) and consisted of the sample sounds of Wizoo acoustic piano. The musical patterns were played on the piano (standard tones) with the exception of the third tone, that was replaced with one of the following six types of deviants: pitch, timbre, localization, intensity, slide or rhythm. Deviants were generated in Adobe Audition (Adobe Systems Incorporated©) by modifying the sound features of interest a follows:

− **Pitch**: the tone was mistuned by 24 cents (downwards tuning for the major mode and upwards tuning for the minor mode).
− **Timbre**: the timbre of the tone was altered with the “old-time radio” effect of Adobe Audition with a 4-channel parametric equalizer.
− **Location**: the amplitude of the right channel was decreased of 10dB, which resulted in a sound perceived as coming from a localization centered to the left (∼70°) compared to the midline.
− **Intensity**: the original intensity was reduced by 6dB.
− Slide: gradual change of the pitch from two semitones below up to the standard over the sound presentation.
− **Rhythm**: 40ms shorter than the standard tone and maintaining the same interstimulus interval (ISI), resulting in the consequent tone arriving earlier than expected.

The musical patterns were played with each of the 24 possible keys (12 major and 12 minor), with the key changing once every six patterns in a pseudo-random order. Each tone (except for the rhythm deviant, which was 160ms long) had a duration of 200ms and 5ms of raise and fall time, with an ISI of 5ms. Each deviant was presented 144 times in pseudo-random order, 50% of which was played in major mode and the other 50% in minor mode. The total duration of the paradigm was 12 minutes. The stimuli were randomized in Matlab and delivered with Presentation software (Neurobehavioral Systems, Berkley, CA). The auditory stimulation was presented through a pair of pneumatic headphones (Sennheiser HD 210) at a comfortable volume level established during the sound-check prior to the actual measurement. Participants were seated in a chair with their heads placed in the helmet-like space of the MEG device. Recordings were taken while the auditory paradigm was presented. Participants passively listened to the sounds while watching a silenced movie of their choice with subtitles.

### Data acquisition

#### Genotyping

DNA analyses were carried out at THL Biobank, National Institute for Health and Welfare, Helsinki, Finland. Deoxyribonucleic acid (DNA) was isolated from blood samples complying with standard extraction protocols. The DNA extraction from K2-EDTA-blood tubes was performed with chemagic 360 instrument and the CMG-704 kit (PerkinElmer), which uses magnetic bead technology. The DNA concentration was measured with Quant-iT™ PicoGreen™ dsDNA Assay Kit after it was eluted in 400µl 10mM Tris-EDTA elution buffer (PerkinElmer). Aliquotes of DNA samples were produced with Tecan Genesis/Tecan Freedom Evo and then shipped on dry ice for genetic analyses. BDNF Val66Met was genotyped with Illumina Infinium PsychArray BeadChip and quality control (QC) was assessed with PLINK. Some markers were removed due to pattern missingness (>5%), low minor allele frequency (< 0.01) and Hardy-Weinberg equilibrium (p-value < 1×10-6). Individual participants were checked for missing genotypes (>5%), relatedness (identical by descent calculation, PI_HAT>0.2) and population stratification (multidimensional scaling).

#### MEG data acquisition

The data were collected with a 306-channel Vectorview whole head MEG scanner (Elekta Neuromag, Elekta Oy, Helsinki, Finland) at the Biomag Laboratory of the Helsinki University Central Hospital. The recordings were carried out in an electrically and magnetically shielded room (ETS-Lindgren Euroshield, Eura, Finland). The MEG device was comprised of 102 axial magnetometers and 102 pairs of planar gradiometers, for a total of 306 SQUID sensors. MEG data were recorded with a sampling rate of 600Hz.

#### MRI data acquisition

We acquired MRI data employing a 3T MAGNETOM Skyra whole□body scanner (Siemens Healthcare, Erlangen, Germany) and a standard 32□channel head□neck coil. The recordings were carried out at the Advanced Magnetic Imaging (AMI) Centre (Aalto University, Espoo, Finland) on a separate date either before or after the MEG session.

T1□weighted structural images were collected for individual coregistration with MEG data and proper estimation of neural sources with the following acquisition parameters: 176 slices; slice thickness□=□1 mm; field of view□=□256 × 256 mm; interslice skip□=□0 mm; matrix□=□256 × 256; pulse sequence□=□MPRAGE.

### Data preprocessing

MEG sensor data of both planar gradiometers (n=204) and magnetometers (n=102) were pre-processed with MaxFilter 2.2 (Taulu & Simola, 2006). Interference originated from external and nearby sources was attenuated by applying signal space separation and the signal for head movement was adjusted. The sampling rate was 600Hz and no down-sampling was performed. The preprocessed data were converted into SPM objects (Penny et al., 2007) and further analyzed in Matlab (Math-Works, Natick, Massachusetts, United States of America), with OSL, a free source toolbox using a combination of Fieltrip (Oostenveld et al., 2011), SPM (Penny et al., 2007) and FSL (Woolrich et al., 2009) functions, together with in-house-built functions. The data was then low-pass filtered at 30Hz. Minimum parts of the signal contained bad trials that were removed manually following visual inspection. Heartbeat and eye blink related artefacts were detected by means of independent component analysis (ICA) and removed manually. The signal was epoched into 500ms segments (from -100ms to 400ms) and baseline corrected at -100ms. Difference waveforms were created by subtracting the average standard waveform from each of the deviant waveforms, as widely done in MEEG studies on MMN (Alho, 1995; Näätänen et al., 2007).

### BDNF and neural responses to deviants

#### MEG sensors

First, an average of the trials was computed independently for each deviant and each couple of planar gradiometers was combined by mean root square. Subsequent statistical analyses were performed on gradiometers only because of their better signal-to-noise ratio compared to magnetometers (in Bonetti et al. 2017, 2018, and Haumann et al., 2016 quantitative measures of signal-to-noise ratio for this same dataset are shown). Such analyses were performed in Matlab using a combination of built-in and in-house-built functions and scripts. Second, an ANOVA was used to compare the neural responses to deviants in the four experimental groups (M1, M0, NM1, NM0). This analysis was computed for each planar gradiometer and each time-point within the time-range 0 – 400ms and then corrected for multiple comparisons using the Tukey-Kramer post-hoc. Thus, p-values were considered significant only when both the ANOVA and the Tukey-Kramer post-hoc were significant. Then, such p-values were binarized by setting 1 when the p-value was significant (adjusted α = .05) and 0 when it was not significant. The binary data were then rearranged in a 3D matrix that recreated a 2D approximation of the MEG channels layout for each time point.

To further correct for multiple comparisons, clusters of 1s were identified and their significance assessed by running Monte-Carlo simulations. To do so, first we calculated the cluster sizes of the original binary matrix. Then, we computed 1000 permutations of such matrix and identified the maximum cluster size of 1s for each permutation. This procedure allowed us to build the distribution of maximum cluster size of the permuted data. Finally, original clusters with a size bigger than 99.9% of permuted data were considered as significant. MCS was used to run two complementary analyses: first, to assess when the activity of M1 was greater than the other experimental groups, as stated by our hypothesis; then, to evaluate when the activity of M1 was smaller than the other experimental groups, as a control.

### Source reconstruction

#### Beamforming

The sources of the brain activity measured by MEG were reconstructed in source space by means of an overlapping-spheres (forward model) approach and a beamformer approach (inverse model) (Hillebrand and Barnes, 2005). Signal from both magnetometers and gradiometers and an 8-mm grid were used. The forward model depicted the MNI-co-registered anatomy (obtained through MRI structural T1 recording for each participant) as a simplified geometric volume by using a set of spheres (Huang et al., 1999). The inverse model sequentially applied a set of weights to the source locations to isolate the contribution of each source to the general MEG activity recorded by the sensors at each time point (Brookes et al., 2007; Hillebrand and Barnes, 2005).

#### General Linear Model (GLM) and cluster-based permutation test

For each experimental condition, and independent GLM was calculated sequentially for each time point at each dipole location (Hunt et al., 2012). The first-level analysis consisted of calculating contrasts between the brain activity recorded for the six deviants and the standard independently for each participant. These results were then submitted to two group-level analysis. The first one did not differentiate between different groups of participants, but merely assessed the quality of the source reconstruction (**Figure 1C**). The second analysis used paired-sample t-tests to contrast M1 vs M0. Here, a spatially smoothed variance computed with a Gaussian kernel (full-width at half-maximum: 50 mm) was employed. Finally, to correct for multiple comparisons, a cluster-based permutation test (Hunt et al., 2012) with 5000 permutations was performed on the second-level analysis. In this case, we analysed only the significant time-windows emerged from the MEG sensor level analysis independently for each deviant, with an α level = .05, which corresponds to a cluster forming threshold t-value = 1.8.

## Acknowledgements

We are glad to thank the team who contributed to the “Tunteet” neuroscientific and psychological data collection: Satu Palva, Johanna Normström, Brigitte Bogert, Benjamin Gold, Mikko Heimölä, Taru Numminen-Kontti, Toni Auranen, Jyrki Mäkelä, Marita Kattelus, Mikko Sams, Mari Tervaniemi, Petri Toivainen, Anja Thiede, David Ellison, Chao Liu, Alessio Falco, Suvi Lehto and Simo Monto.

## Authors contributions

S.E.P. Bruzzone: Conceptualization, data analysis, writing of the original draft, editing and review of the manuscript, figures design and execution.

L. Bonetti: Conceptualization, data analysis, editing and review of the manuscript, figures design.

M. Kliuchko: Study design, data acquisition, review of the manuscript.

T. Paunio: Genotyping, data acquisition, review of the manuscript.

K. Kantojärvi: Genotyping, data acquisition, review of the manuscript.

P. Vuust: Editing and review of the manuscript, supervision, funding.

E. Brattico: Conceptualization, data acquisition, study design, editing and review of the manuscript, supervision, funding.

## Data availability statement

The code and anonymized neuroimaging data from the experiment are available upon reasonable request.

## Supplementary material

**Table ST1** – Neural sources generating the MMN responses. Brain region, hemisphere, t-values and spatial coordinates describing the significant brain sources of MMN for each deviant.

**Table ST2** – MEG sensor results and neural sources obtained by contrasting the experimental groups of musicians (M1 and M0). The brain activity reported here as significant is based on the results from the post-hoc comparisons obtained from the ANOVA performed on each time point. For each deviant, the time window (starting at Time 1 and ending at Time 2) and the sensors in which significant brain activity was detected are reported.

